# Conserved noncoding elements influence the transposable element landscape in *Drosophila*

**DOI:** 10.1101/257907

**Authors:** Manee M. Manee, John Jackson, Casey M. Bergman

**Affiliations:** Faculty of Life Sciences, University of Manchester, Manchester, UK; Current affiliation: National Center for Genomic Technology, King Abdulaziz City for Science and Technology, Riyadh, Saudi Arabia.; Current affiliation: Center of Excellence for Genomics (CEG), King Abdulaziz City for Science and Technology, Riyadh, Saudi Arabia.; Current affiliation: Department of Animal and Plant Sciences, University of Sheffield, Sheffield, UK; Current affiliation: Department of Genetics, University of Georgia, Athens, GA, USA.; Current affiliation: Institute of Bioinformatics, University of Georgia, Athens, GA, USA.

**Keywords:** Noncoding DNA, Conserved Noncoding Elements, Purifying Selection, Transposable Elements, Drosophila

## Abstract

Highly conserved noncoding elements (CNEs) comprise a significant proportion of the genomes of multicellular eukaryotes. The function of most CNEs remains elusive, but growing evidence indicates they are under some form of purifying selection. Noncoding regions in many species also harbor large numbers of transposable element (TE) insertions, which are typically lineage specific and depleted in exons because of their deleterious effects on gene function or expression. However, it is currently unknown whether the landscape of TE insertions in noncoding regions is random or influenced by purifying selection on CNEs. Here we combine comparative and population genomic data in *Drosophila melanogaster* to show that abundance of TE insertions in intronic and intergenic CNEs is reduced relative to random expectation, supporting the idea that selective constraints on CNEs eliminate a proportion of TE insertions in noncoding regions. However, we find no difference in the allele frequency spectra for polymorphic TE insertions in CNEs versus those in unconstrained spacer regions, suggesting that the distribution of fitness effects acting on observable TE insertions is similar across different functional compartments in noncoding DNA. Our results provide evidence that selective constraints on CNEs contribute to shaping the landscape of TE insertion in eukaryotic genomes, and provide further evidence supporting the conclusion that CNEs are indeed functionally constrained and not simply mutational cold spots.

## Introduction

Transposable elements (TEs) are mobile DNA sequences that comprise a significant fraction of the genomes of many multicellular organisms (Elliott and Gregory, 2015), including the model insect species, *Drosophila melanogaster* (Bergman *et al.*, 2006; Sackton *et al.*, 2009). TEs are powerful mutagenic agents that can affect gene expression and genome stability and are responsible for the majority of spontaneous mutations in *D. melanogaster* (Ashburner *et al.*, 2005). While many gaps remain in our understanding of the mechanisms that control TE content in natural populations of *D. melanogaster*, it is well established that TE insertions in the *D. melanogaster* genome are largely restricted to non-coding DNA (reviewed in Barron *et al.* (2014)). Early restriction mapping studies on a limited number of loci revealed that large DNA insertions (assumed to be TEs) were rarely found in transcribed regions (Aquadro *et al.*, 1986; Langley and Aquadro, 1987; Schaeffer *et al.*, 1988; Langley *et al.*, 1988; Aquadro *et al.*, 1992). Subsequent analysis of the *D. melanogaster* reference genome showed that the paucity of TEs in transcribed regions is primarily driven by a strong depletion of the number of TE insertions in exons combined with a weaker reduction in introns (Kaminker *et al.*, 2002; Lipatov *et al.*, 2005). More recently, analysis of population genomic data has confirmed that TE insertions are rare in *D. melanogaster* exonic regions (Kofler *et al.*, 2012; Cridland *et al.*, 2013; Zhuang *et al.*, 2014).

The under-representation of TEs in *D. melanogaster* exons is most likely explained by natural selection purging TE insertions that disrupt gene function from natural populations (Lipatov *et al.*, 2005; Petrov *et al.*, 2011; Kofler *et al.*, 2012). In general, TE insertions in *D. melanogaster* are thought to be under some form of purifying selection, based on the observation that they typically have lower allele frequencies relative to single nucleotide polymorphisms (SNPs) from the same population (Aquadro *et al.*, 1986; Langley and Aquadro, 1987; Schaeffer *et al.*, 1988; Langley *et al.*, 1988; Aquadro *et al.*, 1992; Cridland *et al.*, 2013). However, few studies have directly investigated the allele frequency distribution of TE insertions in exons, principally because of the lack of data, and past studies have led to mixed conclusions. Analysis of a small sample of exonic TE insertions using a pool-PCR strategy suggested their allele frequencies did not differ substantially from non-exonic TE insertions with similar genomic properties (Lipatov *et al.*, 2005). In contrast, genome-wide analysis using pool-seq data showed a reduction in median allele frequencies for TE insertions in exons relative those found in intergenic regions (Kofler *et al.*, 2012).

In addition to effects manifest at the RNA or protein level, it is also possible TE insertions may be selected for their effects at the DNA level in noncoding regions, for example by interfering with *cis*-regulatory elements (Geyer *et al.*, 1990; Lerman and Feder, 2005). While comprehensive *cis*-regulatory maps for *D. melanogaster* remain incomplete (Negre *et al.*, 2011; Arnold *et al.*, 2013), it is well established that highly conserved noncoding elements (CNEs) are an abundant component of the *D. melanogaster* genome (Bergman and Kreitman, 2001; Siepel *et al.*, 2005) and that CNEs often overlap with known *cis*-regulatory elements (Emberly *et al.*, 2003; Brody *et al.*, 2012). It has been estimated that 30%-40% of sites in *D. melanogaster* noncoding DNA are contained in CNEs (Siepel *et al.*, 2005), and population genetic analysis has shown that these CNEs are maintained by purifying selection (Casillas *et al.*, 2007). Thus, CNEs represent an abundant class of noncoding features under purifying selection that may influence the landscape of TE insertions. Previous work showed that artificially-induced TE insertions are depleted in the most highly conserved CNEs (so-called “ultra-conserved elements”) (Makunin *et al.*, 2013). However the non-random target preferences, requirement for marker gene activation in TE detection, and experimental origin of the TEs analyzed by Makunin *et al.* (2013) do not allow conclusions to be drawn about CNE-based constraints on insertion of the endogenous set of TE families in natural populations. Resolving whether CNEs influence the landscape of TE insertion in natural populations of *D. melanogaster* will provide further insight into the factors governing TE dynamics in this species, and contribute to our broader understanding of the forces that shape genome organization and molecular evolution in general.

Here we use genome-wide datasets of “non-reference” TE insertions (i.e. TEs identified in a resequenced sample that are not present in the reference genome) from a North American population of *D. melanogaster* (Mackay *et al.*, 2012; Lin-heiro and Bergman, 2012; Zhuang *et al.*, 2014) to investigate whether selective constraints on CNEs influence the landscape of TE insertions in noncoding DNA. These datasets allow unprecedented insight into this fundamental question by providing large samples of naturally-occurring TE insertions mapped at nucleotidelevel resolution in individual strains of *D. melanogaster*. We initially establish that signals of purifying selection can be observed in our data by confirming past results that the abundance of TE insertions is strongly reduced in exonic regions and weakly reduced in intronic regions relative to intergenic regions. We then show that the abundance of TE insertions is significantly reduced in both intronic and intergenic CNEs relative random expectations. In contrast to the clear signals of purifying selection on TE abundance, we find that the derived allele frequency (DAF) spectrum for TE insertions inferred from strain-specific genome sequences does not vary significantly across different functional compartments of the *D. melanogaster* genome. Our results provide systematic evidence that selective constraints on CNEs in noncoding regions influence the landscape of TE insertion in *D. melanogaster*. However, the proportion of TE insertions we estimate to be eliminated from CNEs is lower than in exonic regions, suggesting that many noncoding functional elements to harbor viable TE insertion mutations in natural populations of *D. melanogaster*. Our results also suggest that the evolutionary forces governing the abundance of TE insertions in different functional compartments of the *D. melanogaster* genome may be decoupled from those controlling the allele frequency of observable TE insertions in natural populations.

## Materials and Methods

### Data Sets

Annotations of genes (flyBaseGene), TEs in the reference genome (rmsk), and conserved elements (phastCons15way) on Release 5 (dm3) coordinates of the *D. melanogaster* genome were obtained from UCSC Genome Browser (Siepel *et al.*, 2005; Smit *et al.*, 2013; Gramates *et al.*, 2017; Tyner *et al.*, 2017). Annotations of non-reference TE insertion in the *Drosophila* Genetic Reference Panel (DGRP) of *D. melanogaster* strains from Raleigh, NC (Mackay *et al.*, 2012) were obtained from supplementary materials of papers describing two different TE detection methods: `ngs_te_mapper` (Linheiro and Bergman, 2012) and `TEMP` (Zhuang *et al.*, 2014).

The `ngs_te_mapper` dataset consists of non-reference TE insertions from 37 long terminal repeat (LTR) retrotransposon and terminal inverted repeat (TIR) transposon families on the major chromosome arms (chrX, chr2L, chr2R, chr3L, chr3R and chr4) identified using whole-genome Illumina shotgun sequence data in 166 DGRP strains (Linheiro and Bergman, 2012). A new BED file for this dataset was generated by Dr. Raquel Linheiro (personal communication) that encodes the number of DGRP strains in which each insertion was found in the score column (Linheiro and Bergman, 2014). The `TEMP` dataset consists of non-reference TE insertions from 56 LTR retrotransposon, non-LTR retrotransposon and TIR transposon families identified using whole-genome Illumina shotgun sequence data in 53 DGRP strains (Zhuang *et al.*, 2014). We transformed the original `TEMP` dataset from from https://zlab.umassmed.edu/TEMP/TEMP_resources/DGRP_53lines_TE_polymorphisms.tar.gz to match the format of the ngs_te_mapper dataset as follows. TE insertions in *.insertion.refined.bp.refsup files were merged across all strains, then insertions supported by split-read data on both ends of the TE found on the major chromosome arms (chrX, chr2L, chr2R, chr3L, chr3R and chr4) were extracted, converted to BED format, sorted, and clustered using BEDtools complement (-s -d 0) (Quinlan and Hall, 2010). The number of strains per cluster containing a TE insertion for the same TE family on the same strand was then encoded in the score column of a BED-formatted file. For both datasets, a small number of TE insertions were predicted to occur at the same location, either from closely related TE families (e.g. *Stalker vs. Stalker 4*) or for TIR elements predicted on opposite strands at the same location (e.g. *S* element). We kept one of these redundant annotations based on the first occurrence in the dataset. Finally, we excluded all *P* element insertions from both datasets, since this TE family is known to have a strong non-random preference to insert around transcriptional start sites (Spradling *et al.*, 1995; Bellen *et al.*, 2004; Kofler *et al.*, 2015).

### Assigning TE insertions to genomic compartments

We partitioned regions of the *D. melanogaster* genome into mutually-exclusive exonic, intronic and intergenic compartments based on the gene structures in the dm3 flyBaseGene track using the overlapSelect and BEDtools intersect, complement, subtract tools (Kuhn *et al.*, 2013; Quinlan and Hall, 2010). Each tool was run using default parameter settings. Our partitioning strategy follows Lipatov *et al.* (2005) and assumes a hierarchy of functional constraints for genomic regions that have multiple annotation states due to alternative splicing or promoter usage: namely, functional constraints on exonic regions take precedence over intronic regions, and constraints on intronic regions take precedence over intergenic regions. Exonic regions span the union of all exon intervals in the genome and include both coding sequences (CDS) and untranslated regions (UTRs). Intronic regions were defined as the complement of exonic regions in genomic intervals spanned by at least one transcript model. Intergenic regions were defined as the complement of all exonic and intronic regions. Intronic and intergenic regions were further partitioned into CNEs and spacers using the dm3 phastCons15way track. Spacers are defined as the noncoding regions complementary to CNEs that exhibit low primary sequence conservation (Bergman *et al.*, 2002; Casillas *et al.*, 2007). Reference TE intervals were subtracted from all exonic, intronic, intergenic, CNE, and spacer compartments.

Non-reference TE insertions in the `ngs_te_mapper` and `TEMP` datasets were then assigned to genomic compartments in high recombination regions using overlapSelect (Kuhn *et al.*, 2013). The locations of the non-reference TE insertions studied here are annotated as their target site duplication (TSD) (Linheiro and Bergman, 2012; Zhuang *et al.*, 2014), which span small intervals (typically <10 bp) on reference genome coordinates and can therefore overlap the boundaries of neighboring genomic compartments. To avoid counting TEs that overlap boundaries multiply or partially in different compartments, a series of filtering steps was implemented to identify TE insertions that overlap intronic/exonic, intergenic/exonic and CNE/spacer boundaries. Each distinct category of “overlapping” TE insertions is mutually exclusive with other overlapping or “pure” compartments. Nonreference TEs found fully or partially in annotated in reference TE intervals were removed from all datasets.

We restricted our analysis to regions of the *D. melanogaster* Release 5 genome sequence with normal rates of recombination using criteria established in previous population genomic analyses of TEs in *D. melanogaster* (Cridland *et al.*, 2013, 2015): chrX:300000-20800000, chr2L:200000-20100000, chr2R:2300000-21000000, chr3L:100000-21900000, chr3R:600000-27800000. Low recombination regions were excluded because of the high density of reference TE insertions in these regions (Bartolome *et al.*, 2002; Bergman *et al.*, 2006), which poses challenges to identifying non-reference TE insertions as well as defining CNEs using comparative genomic data. Furthermore, the efficacy of natural selection on individual alleles is reduced in regions of the *Drosophila* genome with low rates of recombination because of the confounding effects of selection on linked sites extending over larger regions (Presgraves, 2005; Haddrill *et al.*, 2007). Normally-recombining regions occupy 89.8% of the 120 Mb Release 5 genome. The numbers of nucleotides and proportion of the genome spanned by each compartment in normally-recombining and normally-recombining noncoding regions are shown in Table. 1 and 2, respectively. The majority of non-reference TEs in both datasets studied here were located in normally-recombining regions (`ngs_te_mapper`: n = 6099/6747, 90.4%; `TEMP`: 4688/5331, 87.9%).

**Table 1:** TE insertions in normal recombination regions. Columns contain the coverage (in base pairs) and percent of the normally-recombining genome covered for exonic, intronic and intergenic regions followed by the number and percent of TE insertions found fully in exonic, intronic and intergenic regions or spanning intron/exon and intergenic/exon boundaries for both `ngs_te_mapper` and `TEMP`. Overlap categories have “n.a.” for coverage and percent of the normally-recombining genome covered since boundaries between compartments do not occupy any space. Regions of the reference genome identified by RepeatMasker as TE were excluded from all other compartments and any non-reference TE in these regions are included in the “Reference TE” compartment. Regions of normal recombination were defined by Cridland *et al.* (2013).

| Region | Coverage (bp) | % normal rec.<br>genome | # `ngs_te_mapper` TE | % `ngs_te_mapper` TE | # `TEMP` TE | % `TEMP` TE |
| --- | --- | --- | --- | --- | --- | --- |
| Exon | 27502613 | 25.4 | 399 | 6.5 | 278 | 5.9 |
| Intron | 38960671 | 36 | 2743 | 45 | 2153 | 45.9 |
| Intron/Exon | n.a. | n.a. | 5 | 0.1 | 7 | 0.1 |
| Intergenic | 37804929 | 35 | 2905 | 47.6 | 2210 | 47.1 |
| Intergenic/Exon | n.a. | n.a. | 9 | 0.1 | 4 | 0.1 |
| Reference TE | 3831787 | 3.5 | 38 | 0.6 | 36 | 0.8 |
| Total | 108100000 | 100 | 6099 | 100 | 4688 | 100 |

**Table 2:** TE insertions in noncoding regions with normal recombination Columns contain the coverage (in base pairs) and percent of the normally-recombining noncoding genome covered by CNEs and spacers for introns and intergenic regions followed by the number and percent of TE insertions found fully in CNEs and spacers or spanning CNE/spacer boundaries for both `ngs_te_mapper` and `TEMP`. Overlap categories have “n.a.” for coverage and percent of the normally-recombining noncoding genome covered since boundaries between compartments do not occupy any space. Regions of the reference genome identified by RepeatMasker as TE and any non-reference TE in these regions were excluded from all compartments. Regions of normal recombination were defined by Cridland *et al.* (2013).

| Region | Coverage (bp) | % normal rec.<br>noncoding genome | # `ngs_te_mapper` TE | % `ngs_te_mapper` TE | # `TEMP` TE | % `TEMP` TE |
| --- | --- | --- | --- | --- | --- | --- |
| Intronic CNE | 14093340 | 18.4 | 747 | 13.2 | 500 | 11.5 |
| Intronic spacer | 24867331 | 32.4 | 1842 | 32.6 | 1458 | 33.4 |
| Intronic CNE/Spacer | n.a. | n.a. | 154 | 2.7 | 195 | 4.5 |
| Intergenic CNE | 14749396 | 19.2 | 813 | 14.4 | 577 | 13.2 |
| Intergenic spacer | 23055533 | 30 | 1928 | 34.1 | 1447 | 33.2 |
| Intergenic CNE/Spacer | n.a. | n.a. | 164 | 2.9 | 186 | 4.3 |
| Total | 76765600 | 100 | 5648 | 100 | 4363 | 100 |

### Testing for purifying selection on TE insertions

We tested for depletion of TE insertions in different genomic compartments relative to random expectations using a permutation approach, which accounts for the empirical length distributions of intervals in different genomic compartments and accommodates the variable lengths of TSDs for non-reference TEs. Random TE insertion was simulated using BEDTools shuffle to permute the location of TE insertions in different compartments of the Release 5 genome. Random TE insertions were required to be placed within their same chromosome *(-chrom* option), were not allowed to overlap each other *(-noOverlapping* option), and were not allowed to land in regions of the reference genome annotated as TE by Repeat-Masker (Smit *et al.*, 2013). We attempted to control for the effects of selection on non-focal genomic compartments by excluding TEs from these regions and blacklisting insertion in non-focal regions using the BEDtools shuffle *-excl* option. The *-seed* option was used to allow results of each run to be replicated. TE insertions in randomized datasets were then assigned to genomic compartments as described above.

A series of permutation tests were performed to test the null hypothesis of random TE insertion across various sets of genomic compartments. All permutation tests were restricted to normally-recombining regions of the genome as defined above. First, TE insertions observed in all compartments were allowed to randomly insert into all compartments to test if TEs are depleted in exonic regions relative to noncoding DNA. Second, TE insertions observed in noncoding regions were allowed to randomly insert in noncoding regions to test if TEs are depleted in introns relative to intergenic regions, independent of the effects of purifying selection on exonic regions. Third, TE insertions observed in intronic regions were allowed to randomly insert in intronic regions to test if TEs are depleted in intronic CNEs relative to intronic spacers, independent of the effects of purifying selection on exonic or intergenic regions but accounting for potential selection on introns. Finally, TE insertions observed in intergenic regions were allowed to randomly insert in intergenic regions to test if TEs are depleted in integenic CNEs relative to intergenic spacers, independent of the effects of purifying selection on exonic or intronic regions. For each test, 10,000 permutations were performed to provide a distribution of outcomes under the null hypothesis of random insertion.

Additionally, we tested whether the derived allele frequency (DAF) of TE insertions in putatively selected genomic compartments (exonic regions, CNEs) differed from control regions (intergenic spacers). Following previous efforts testing whether CNEs are cold spots of point mutation (Drake *et al.*, 2006; Casillas *et al.*, 2007), the null hypothesis of no difference in DAF between “selected” and “control” compartments was tested using a non-parametric Wilcoxon rank sum test. DAF tests of TE insertion allele frequencies in CNEs *vs.* spacers were performed separately for intronic and intergenic regions. As in related work (Petrov *et al.*, 2011; Kofler *et al.*, 2012; Cridland *et al.*, 2013), we assumed all TE insertions represent the derived state since, with the exception of the *INE-1* family that is not studied here (Singh *et al.*, 2005; Wang *et al.*, 2007), few TE insertions in *D. melanogaster* are thought to have occurred prior to speciation (Caspi and Pachter, 2006; Bergman and Bensasson, 2007; Sackton *et al.*, 2009). Rare TE insertions spanning intron/exon on intergenic/exon boundaries were excluded from DAF analysis because of their low sample sizes. However, TE insertions spanning CNE/spacer boundaries were relatively common, and thus were analyzed as distinct class and compared to TEs contained fully within spacers.

All graphical and statistical analyses were performed in the R programming environment (version 3.4.0) (R Core Team, 2016).

## Results

### TE insertions are depleted in conserved noncoding elements

To understand whether selective constraints on noncoding DNA influence patterns of TE insertion, we analyzed the abundance of non-reference TEs insertions in different functional genomic compartments of the *D. melanogaster* genome. We first assigned non-reference TE insertions in normally-recombining regions to functional compartments based on gene and conserved element annotations (see Materials and Methods for details). We then tested for depletion of non-reference TE insertions in genomic regions with putatively higher levels of functional constraint (i.e. exonic regions, CNEs) by comparing observed numbers of TEs in these regions to an empirical null distribution based of 10,000 random permutations of the observed TE insertion datasets. Recent studies have shown that no single bioinformatic system can comprehensively identify all non-reference TE insertions in resequencing data (Nelson *et al.*, 2017; Rishishwar *et al.*, 2017). Therefore, we used two independent non-reference TE insertion datasets, `ngs_te_mapper` (Linheiro and Bergman, 2012) and `TEMP` (Zhuang *et al.*, 2014), both derived from the same sample of strain-specific genome sequences isolated from a North American population of *D. melanogaster* (Mackay *et al.*, 2012). Both datasets analyzed here both provide large samples of non-reference TE insertions with nucleotide-level resolution based on split-read information, which improves identification of allelic insertions occupying the same insertion site in different strains and assignment of TE insertion sites to specific genomic compartments.

As a positive control, we first tested whether the previously-reported depletion of TE insertions in *D. melanogaster* exonic regions (Lipatov *et al.*, 2005; Kofler *et al.*, 2012; Cridland *et al.*, 2013) could be observed in the `ngs_te_mapper` and `TEMP` datasets using our randomization procedure. As shown in Table 1, several hundred TE insertions in exonic regions can be found in natural populations of *D. melanogaster* (see also Kofler *et al.* (2012); Cridland *et al.* (2013)). Nevertheless, we observed a clear depletion of TE insertions in exonic regions relative to random expectations (Figure 1A), coupled with a concomitant excess in intronic regions (Figure 1B) and intergenic regions (Figure 1C). We estimate a 4-fold *(P <* 1e — 04) and 4.35-fold (P < 1e — 04) reduction in TEs in exonic regions relative to the median of random outcomes for the `ngs_te_mapper` and `TEMP` datasets, respectively (Figure 1A). We also detected evidence for a significant depletion of TE insertions spanning intron/exon boundaries (Figure 1D) for both `ngs_te_mapper` (4.6-fold reduction, P = 1e — 04) and `TEMP` (5.9-fold reduction, P < 1e — 04), consistent with the presence of “hazardous zones” for TE insertion near intron-exon junctions shown previously in humans (Zhang *et al.*, 2011). In contrast, we observed no significant depletion of TEs at intergenic/exon boundaries (Figure 1E; `ngs_te_mapper`: P = 0.98; `TEMP`: P = 0.27). These results support previous analyses that TEs are selectively eliminated from exonic regions (Lipatov *et al.*, 2005; Petrov *et al.*, 2011; Kofler *et al.*, 2012; Cridland *et al.*, 2013), and demonstrate that our approach can detect selective constraints on TE insertions that are assumed to exist in the *D. melanogaster* genome.

**Figure 1:**
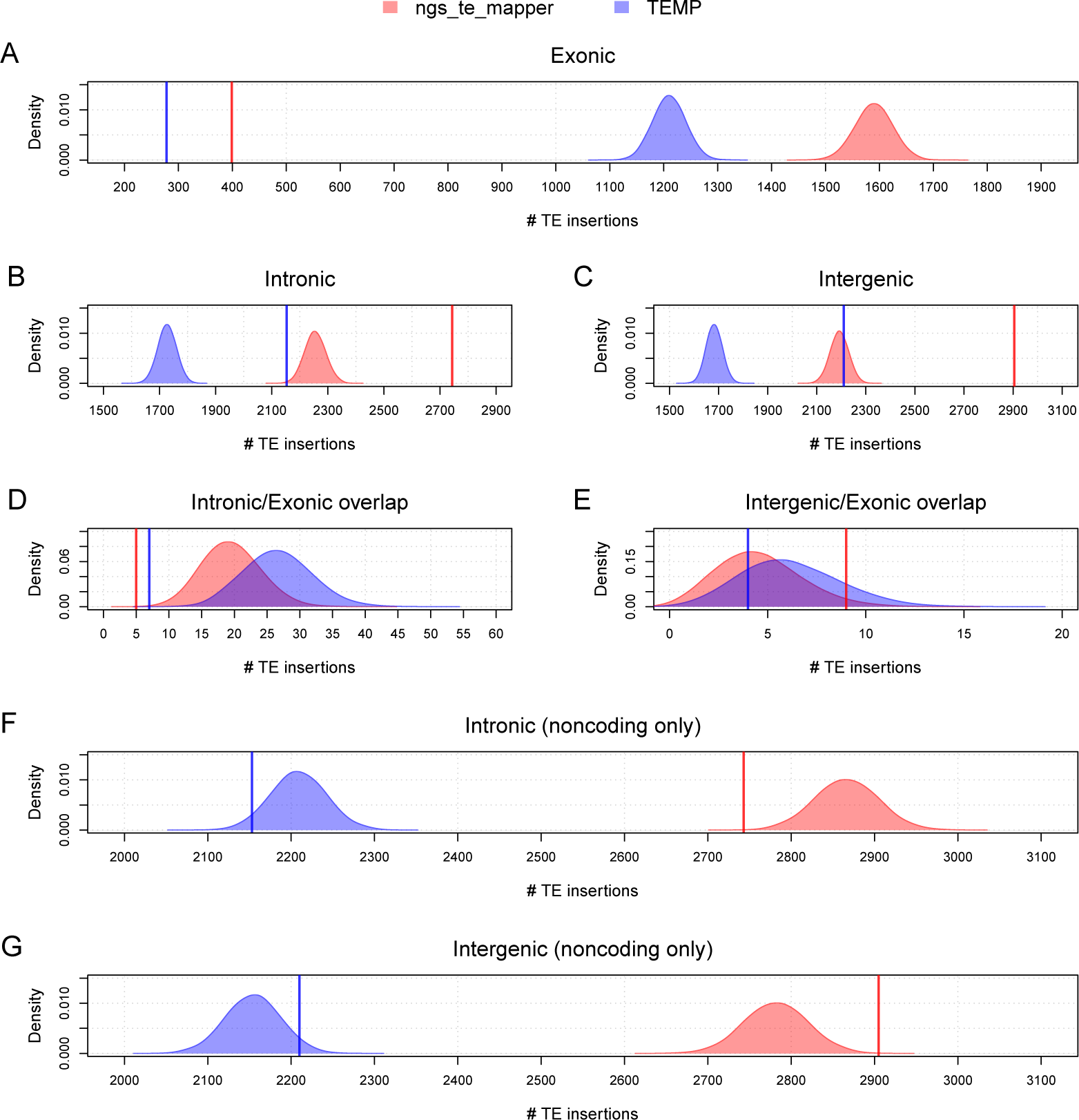
TEs in normally-recombining regions of the *D. melanogaster* genome are depleted in exonic and intronic regions. Observed numbers of TEs in different genomic compartments are shown as vertical lines for `ngs_te_mapper` (red) and `TEMP` (blue). Empirical null distributions of the numbers of TEs in different genomic compartments in 10,000 random permutations are shown as density plots for `ngs_te_mapper` (red) and `TEMP` (blue). All permutation analyses were restricted to normally-recombining regions of the *D. melanogaster* genome as defined by Cridland *et al.* (2013). Permutation analyses were conducted across all compartments (A-E), or in noncoding regions only (F,G). Observed and simulated numbers of TEs were counted in exonic regions (A), intronic regions (B,F), intergenic regions (C,G), intronic/exonic boundaries (D), and intergenic/exonic boundaries (E). Observed TEs overlapping intron/exon boundaries or intergenic/exon boundaries were excluded from permutation analyses in noncoding regions only (F,G). Regions of the reference genome identified by RepeatMasker as TE sequence and any non-reference TE in these regions were also excluded from all permutation analyses.

We next investigated whether our data provide evidence that purifying selection eliminates a higher proportion of TEs in intronic regions relative to intergenic regions, by permuting the locations of TEs in noncoding regions only. We observed a trend towards fewer TE insertions in intronic regions relative to random expectation (Figure 1F) with a corresponding excess in intergenic regions (Figure 1G) in both datasets. The magnitude of this effect was weak but highly significant in the `ngs_te_mapper` dataset (1.05-fold reduction, P = 3e — 04), and of a similar magnitude but less significant in the `TEMP` dataset (1.02-fold reduction, P = 0.05). Our results support those of Kofler *et al.* (2012) who similarly observed a weak but significant reduction in numbers of TE insertions in intronic regions relative to intergenic regions using pool-seq data, but differ from Cridland *et al.* (2013) who observed more TEs in intronic regions relative to intergenic regions using strain-specific genome data. Together, these results suggest that the TE density in *D. melanogaster* intronic regions is weakly reduced relative to random expectations, but that the proportion of TEs eliminated from intronic regions is not sufficiently large for the effect to be reliably identified in all population genomic datasets.

Finally, we tested whether TE insertions were depleted in CNEs relative to spacer regions (Figure 2). For this analysis, we randomized TE insertions separately within intronic regions and within intergenic regions and accounted for TE insertions spanning CNE/spacer boundaries. We identified several hundred TE insertions that exist in CNEs in both intronic and intergenic regions (Table 2). Nonetheless, we found evidence for a significant depletion in the density of TEs in CNEs in both intronic regions (Figure 2A; `ngs_te_mapper`: 1.21-fold reduction, P < 1e — 04; `TEMP`: 1.31-fold reduction, P < 1e — 04) and intergenic regions (Figure 2B; `ngs_te_mapper`: 1.3-fold reduction, P < 1e — 04; `TEMP`: 1.3-fold reduction, P < 1e — 04). We also observed a weaker trend towards fewer TE insertions overlapping CNE/spacer boundaries relative to random expectation in both intronic regions (Figure 2C; `ngs_te_mapper`: 1.18-fold reduction, P = 0.04; `TEMP`: 1.23-fold reduction, P = 0.002) and intergenic regions (Figure 2D; `ngs_te_mapper`: 1.16- fold reduction, P = 0.16; `TEMP`: 1.28-fold reduction, P = 1e — 04). Correspondingly, we also observe that TE insertions in both datasets are over-represented in spacers in both intronic regions (Figure 2E; `ngs_te_mapper`: 1.11-fold excess, P < 1e — 04; `TEMP`: 1.15-fold excess, P < 1e —04) and intergenic regions (Figure 2F; `ngs_te_mapper`: 1.83-fold excess, P < 1e — 04; `TEMP`: 1.17-fold excess, P < 1e — 04). Overall, these results suggest that while some CNEs tolerate disruption by large TE insertions, constraints on CNEs are substantial enough to eliminate enough TE insertions in CNEs to bias the distribution of observed TE insertions towards spacers in noncoding regions of the *D. melanogaster* genome.

**Figure 2:**
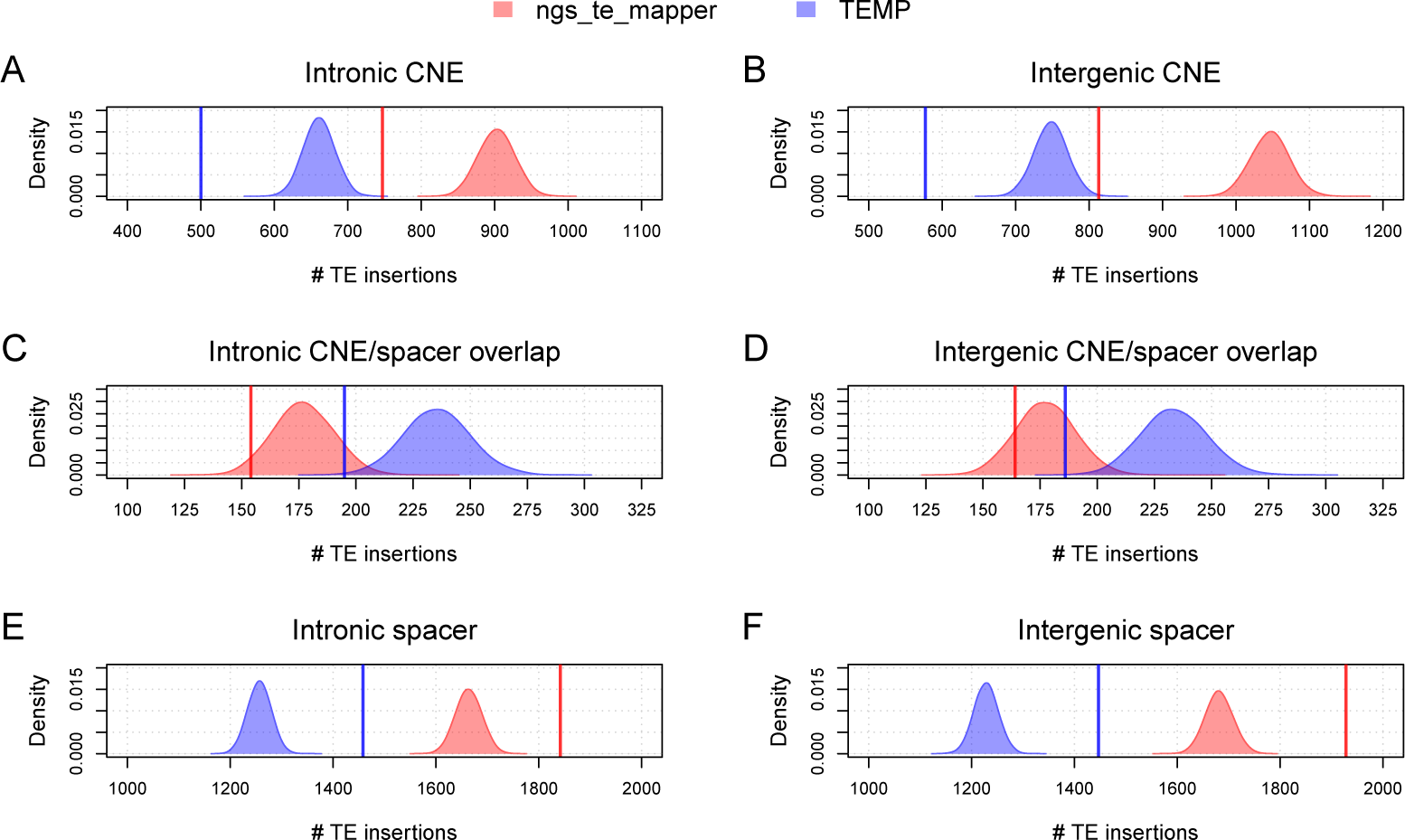
TEs in normally-recombining regions of the *D. melanogaster* genome are depleted in conserved noncoding elements. Observed numbers of TEs in different noncoding compartments are shown as vertical lines for `ngs_te_mapper` (red) and `TEMP` (blue). Empirical null distributions of the numbers of TEs in different noncoding compartments in 10,000 random permutations are shown as density plots for `ngs_te_mapper` (red) and `TEMP` (blue). All permutation analyses were restricted to normally-recombining regions of the *D. melanogaster* genome as defined by Cridland *et al.* (2013). Permutation analyses were conducted across intronic regions only (A,C,E) or intergenic regions only (B,D,F). Observed and simulated numbers of TEs were counted in CNEs (A,B), CNE/spacer boundaries (C,D), or spacers (E,F). The `TEMP` dataset has higher number of observed and expected CNE/spacer overlaps (C,D) despite having fewer TE insertions overall because of a larger average TSD length (7.71 bp) relative to `ngs_te_mapper` (4.73 bp). Observed TEs overlapping intron/exon boundaries or intergenic/exon boundaries were excluded from these analyses. Regions of the reference genome identified by RepeatMasker as TE sequence and any non-reference TE in these regions were also excluded from all permutation analyses.

### Allele frequencies of TE insertions are similar across different functional compartments of the *D. melanogaster* genome

Additional evidence for purifying selection acting to shape the landscape of TE insertions can be obtained from investigating the allele frequencies of TE insertions in population samples. Population genetics theory predicts that natural selection will prevent new deleterious alleles from reaching high population frequency (Fay *et al.*, 2001). If polymorphic TE insertions are weakly negatively selected, they should be skewed towards lower allele frequencies in regions under higher of selective constraint such as exonic regions and CNEs relative to control regions that have weaker functional constraint. A skew in the frequency of *D. melanogaster* SNPs toward rarer alleles has previously been observed in CNEs relative to spacers (Casillas *et al.*, 2007) and in replacement sites relative to silent sites (Huang *et al.*, 2014). However, small indels showed no tendency to be skewed towards rarer alleles in CNEs relative to spacers, suggesting a similar distribution of fitness effects for small indels in both types of noncoding region (Casillas *et al.*, 2007).

Figure 3 shows the DAF spectra for TE insertions in different functional compartments across the *D. melanogaster* genome. Consistent with classical restriction mapping and *in situ* hybridization studies (reviewed in Charlesworth and Langley (1989); Nuzhdin (1999)) and recent strain-specific population genomic data (Cridland *et al.*, 2013), both methods show the expected pattern for TE insertion alleles to be skewed towards rare alleles in all genomic compartments. However, clear differences are observed between `ngs_te_mapper` (Figure 3A) and `TEMP` (Figure 3B) in the overall shape of the DAF spectra across all compartments, with a skew towards more rare alleles in the `ngs_te_mapper` dataset relative to `TEMP`. We interpret overall differences in DAF spectra between TE datasets to result primarily from the higher false negative rate for `ngs_te_mapper` relative to `TEMP` (Nelson *et al.*, 2017) (see see Discussion). Regardless of the cause, comparison of DAF spectra across genomic compartments *within* a dataset should not be substantially compromised, since all compartments are affected by the same methodological biases in TE detection.

**Figure 3:**
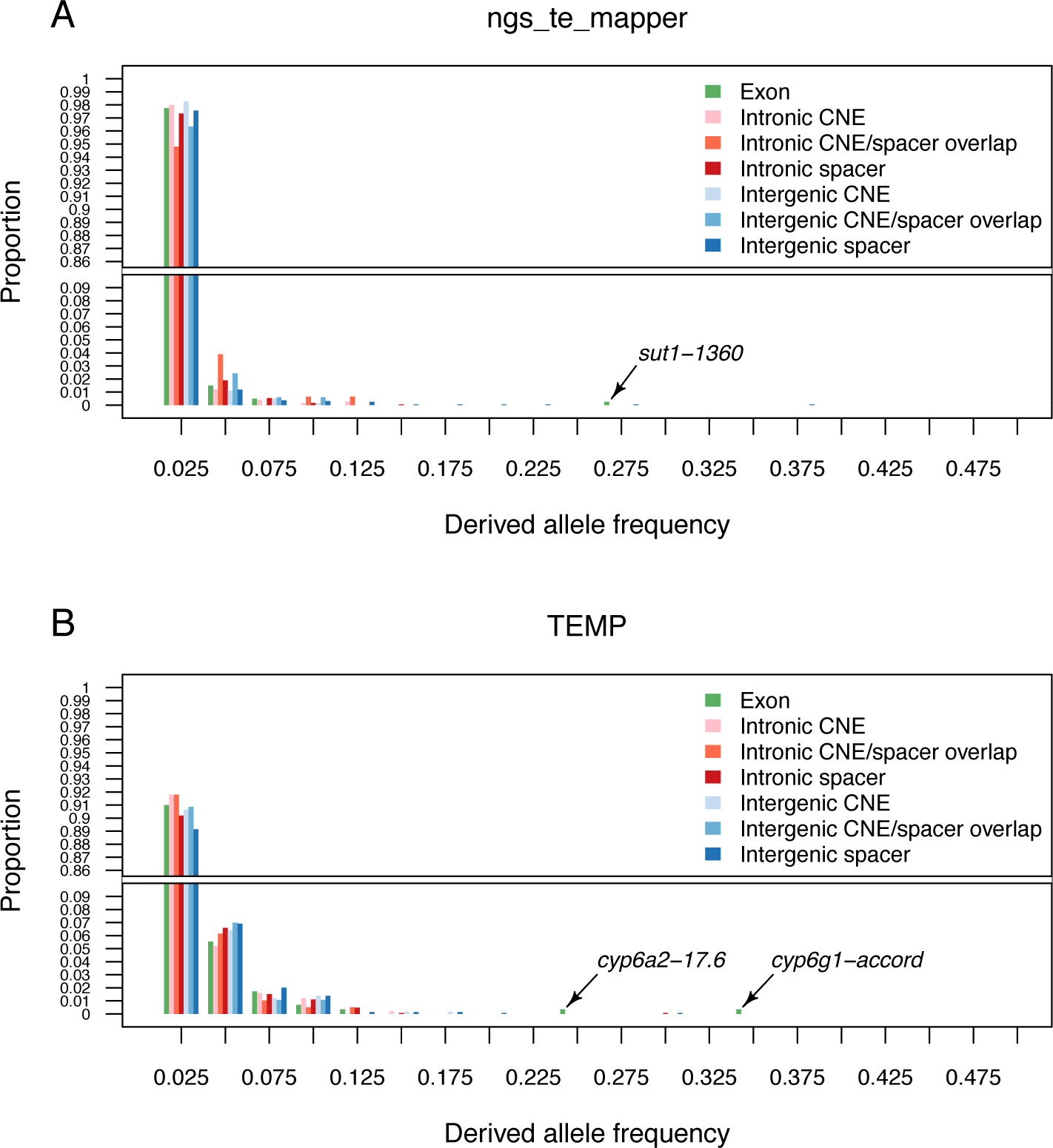
The derived allele frequency (DAF) spectrum for TE insertions is similar across different compartments of the *D. melanogaster* genome. DAF spectra are shown for TE insertions predicted by `ngs_te_mapper` (A) or `TEMP` (B). Allele frequency classes are shown on the X-axis, and the proportion of TE insertions observed in a particular compartment of the genome at that allele frequency is shown on the Y-axis. Note that the Y-axis is split to allow better visualization of the proportion of higher allele frequency classes.

We first performed a control analysis to assess whether the expected skew towards lower allele frequencies could be observed for TE insertion in exonic regions. For this and all subsequent DAF spectra analyses, we used TE insertions in intergenic spacers a control, based on abundance results above showing this compartment was under the weakest selective constraint for TE insertion. As shown in Figure 3, we find no significant differences between the DAF spectra for TEs in exonic regions in either dataset: (ngs_te_mapper: W = 391158.5, P = 0.43; `TEMP`: W = 205299.5, P = 0.36). One possibility for the lack of skew towards rarer alleles for TEs in exonic regions is the presence of a small number of unusually high-frequency exonic TE insertions that are potentially involved in adaptation to insecticide resistance (arrows, Figure 3A,B) (`ngs_te_mapper`: *1360* in *sut1* (Steele *et al.*, 2014); `TEMP`: *17.6* in *cyp6a2* (Waters *et al.*, 1992; Delpuech *et al.*, 1993; Wan *et al.*, 2014), *accord* in *cyp6g1* (Daborn *et al.*, 2002; Chung *et al.*, 2007)). When these putatively-adaptitive outlier loci are excluded, TEs in exonic regions still do not show a consistent skew towards rarer alleles relative to those in intergenic spacers regions: (`ngs_te_mapper`: W = 389232.5, P = 0.5; `TEMP`: W = 203853.5, P = 0.27). These results suggest that the distribution of fitness effects for exonic TE insertions that are not strongly deleterious does not differ substantially from those in intergenic spacers (see also Lipatov *et al.* (2005)).

Next, we tested whether the DAF spectrum for TE insertions in CNEs differed from those in noncoding spacer regions. In this analysis, we also considered the DAF spectrum of TE insertions that spanned CNE/spacer boundaries, because this overlap class is reasonably common and also exhibits a trend towards being depleted in TE insertions (see above). As shown in Figure 3, we found no significant differences in the DAF spectra for TEs in CNEs relative to those in spacer intervals in both intronic regions (`ngs_te_mapper`: *W* = 671827, *P* = 0.19; `TEMP`: *W* = 358690, *P* = 0.29) and intergenic regions (`ngs_te_mapper`: *W* = 767402.5, *P* = 0.2; `TEMP`: *W* = 411058, *P* = 0.31). Likewise, the DAF spectra for TEs overlapping CNE/spacer boundaries did not differ from TEs fully contained in spacer intervals in both intronic regions (`ngs_te_mapper`: *W* = 141937, *P* = 0.98; `TEMP`: *W* = 139781.5, *P* = 0.46) and intergenic regions (`ngs_te_mapper`: *W* = 157028.5, *P* = 0.83; `TEMP`: *W* = 132093, *P* = 0.44). Similar to previous results for small indels (Casillas *et al.*, 2007), these results imply that the distribution of fitness effects on large TE insertions wholly or partially contained in CNEs is not substantially different from that operating on spacer regions in noncoding DNA.

## Discussion

Here we show that the abundance of TE insertions is significantly reduced relative to random expectation in two distinct genomic compartments with known or suspected function: exonic regions and CNEs. In contrast, we find no clear signature for a skew towards lower allele frequencies for TEs in these genomic compartments when compared to regions of the genome under the lowest level of selective constraint. Our results provide the first systematic evidence that selective constraints on CNEs influence the landscape of TE insertion in a eukaryote genome, and provide new evidence supporting the conclusion that CNEs are functionally constrained and not mutational cold spots. Our results also suggest that distribution of fitness effects acting on polymorphic TEs insertions (which have escaped rapid elimination by strong purifying selection) is similar across different functional compartments of the *D. melanogaster* genome.

Our conclusions are derived from two TE insertion datasets (`ngs_te_mapper` and `TEMP`), indicating they are not dependent on the idiosyncracies of a single method for calling TE insertions in short-read resequencing data. Nevertheless, it is important to consider how our results may be affected by the imperfect state of the art in TE calling in terms of positional accuracy and false negative rates (Nelson *et al.*, 2017; Rishishwar *et al.*, 2017). It is unlikely that the depletion of TE insertions we observe is due to imprecise annotation of the TE insertions analyzed here, since under-representation of TEs in exonic regions has been observed previously using a variety of different classical and genomic approaches (Aquadro *et al.*, 1986; Langley and Aquadro, 1987; Schaeffer *et al.*, 1988; Langley *et al.*, 1988; Aquadro *et al.*, 1992; Kaminker *et al.*, 2002; Bartolome *et al.*, 2002; Lipatov *et al.*, 2005; Kofler *et al.*, 2012; Cridland *et al.*, 2013; Zhuang *et al.*, 2014). Likewise, false negatives are unlikely to generate the abundance patterns we observe. For this to be the case, the allele frequency of TE insertions would need to be skewed towards higher frequencies in compartments with lower levels of constraint, so that a higher relative proportion of singleton TE insertion sites would fail to be detected in compartments under higher constraint (leading to an artifactually lower number of insertion sites in high constraint regions). However, we find no evidence for a skew towards higher DAF in compartments with lower levels of constraint in our data (Figure 3).

Although we observe the expected pattern of depletion of TEs in higher constraint regions, we find no difference in the DAF spectra between highly constrained and weakly constrained compartments within either the `ngs_te_mapper` or `TEMP` datasets. It is unlikely that positional inaccuracy or false negatives can explain the lack of difference in the DAF spectra between exonic regions or CNEs and spacers. As above, the high positional accuracy of the `ngs_te_mapper` and `TEMP` datasets mitigates against mis-assignment of TEs to the wrong compartment, which could in principle cause the DAF spectra for different compartments to appear more similar than they really are. Furthermore, in the case of CNEs, we accounted for potential blurring of compartment assignment by showing that the DAF spectra of TEs spanning CNE/spacer boundaries have similar allele frequencies to TEs fully contained within CNEs. Additionally, while it is clear that false negatives distort the DAF spectrum towards rare alleles (Emerson *et al.*, 2008), TEs in our study were detected independent of any information about functional compartment and thus false negatives should affect the DAF spectra for all functional compartments in a similar way.

Importantly, we did observe systematic differences in the DAF across TE detection methods, which has not been discussed sufficiently as an issue in population genomic analysis of TE insertions. Specifically, we find that the DAF for `ngs_te_mapper` is skewed more towards lower frequencies that the DAF for `TEMP` (Figure 3A *vs.* B). We do not interpret this difference among method to result from lower positional accuracy of `ngs_te_mapper` relative to `TEMP` artificially splitting alleles from the same insertion site into several different insertion sites each at lower allele frequency, since both datasets use split-read information. Rather it is more likely this difference in DAF among methods results from the higher false negative rate for `ngs_te_mapper` (58% on simulated data (Nelson *et al.*, 2017)) relative to `TEMP` (10% on simulated data (Nelson *et al.*, 2017)). This observation cautions against naive use of allele frequency data from short-read TE insertion detection methods to test predictions of population genetic models, since the precise shape of the frequency spectrum may be determined by false negative rates of TE detection methods rather than any particular evolutionary force (Emerson *et al.*, 2008). This result also motivates more advanced methods to estimate the TE frequency spectra that incorporate false negative detection rates, similar to methods for estimating the frequency spectrum of SNPs that incorporate false positive rates due to sequencing error (Kim *et al.*, 2011; Nielsen *et al.*, 2012).

Our twin findings of depletion of TEs in functional elements like exonic regions and CNEs coupled with a lack of a skew toward rarer alleles in these regions suggests that the selective mechanism controlling location of TEs in the *D. melanogaster* genome may be decoupled from the forces governing allele frequencies of polymorphic alleles (Petrov *et al.*, 2011). Among competing theories for selective forces acting on TE insertions (Nuzhdin, 1999; Lee and Langley, 2010), it is easiest to interpret the depletion of TEs in exonic regions as being due to the direct effects of TE insertion (Petrov *et al.*, 2011; Kofler *et al.*, 2012) and the same logic should hold for depletion of TEs in CNEs. However, the similarity of DAF spectra in different genomic compartments is consistent with the remainder of TE insertions that are not eliminated from functional elements being governed by a number of evolutionary mechanisms. Polymorphic TE insertions could be at similar allele frequencies in different compartments simply because they inserted at similar distributions of times in the past (Bergman and Bensasson, 2007; Kofler *et al.*, 2012; Blumenstiel *et al.*, 2014). Alternatively, the similar DAF spectra of polymorphic TE insertions in different genomic compartments could reflect similar distributions of selective effects that are independent of the precise location of a TE insertion, which might be expected if the deleterious effects of TE insertion are caused by ectopic exchange events (Petrov *et al.*, 2011; Kofler *et al.*, 2012) or local epigenetic silencing spreading from TE insertions (Lee, 2015; Lee and Karpen, 2017). While our work does not resolve these widely-debated alternatives, it does reveal that the selective effects of TE insertion on conserved elements in noncoding DNA needs to be factored into future models explaining TE evolution in *D. melanogaster* and other species.

## Acknowledgments

The authors would like to thank Raquel Linheiro, Michael Nelson, Florence Gutzwiller and Mar Marzo Llorca for their valuable suggestions throughout this project, and members of the Bergman, Dyer, Hall and White Labs for comments on the manuscript. This work was funded by Life Science and Environment Research Institute, King Abdulaziz City for Science and Technology.

## Author Contributions

CMB conceived and designed the experiments; MMM and JJ carried out the experiments; MMM and CMB analyzed the data; MMM and CMB wrote the manuscript. All authors reviewed the manuscript.

